# Mutations in a set of ancient matrisomal glycoprotein genes across neoplasia predispose to disruption of morphogenetic transduction

**DOI:** 10.1101/2022.06.28.497924

**Authors:** Jimpi Langthasa, Satyarthi Mishra, U Monica, Ronak Kalal, Ramray Bhat

**Author notes:** JL and SM contributed equally to this paper.

## Abstract

Misexpression and remodeling of the extracellular matrix is a canonical hallmark of cancer, although the extent of cancer-associated aberrations in the genes coding for ECM proteins and consequences thereof, are not well understood. In this study, we examined the alterations in core matrisomal genes across a set of nine cancers. These genes, especially the ones encoding for ECM glycoproteins, were observed to be more susceptible to mutations than copy number variations across cancers. We classified the glycoprotein genes based on the ubiquity of their mutations across the nine cancer groups and estimated their evolutionary age using phylostratigraphy. To our surprise, the ECM glycoprotein genes commonly mutated across all cancers were predominantly unicellular in origin, whereas those commonly showing mutations in specific cancers evolved mostly during and after the unicellular-multicellular transition. Pathway annotation for biological interactions revealed that the most pervasively mutated glycoprotein set regulated a larger set of inter-protein interactions and constituted more cohesive interaction networks relative to the cancer-specific mutated set. In addition, ontological prediction revealed the pervasively mutated set to be strongly enriched for basement membrane dynamics. Our results suggest that ancient unicellular-origin ECM glycoproteins were canalized into playing critical tissue morphogenetic roles, and when disrupted through matrisomal gene mutations, associate with neoplastic transformation of a wide set of human tissues.

## Introduction

Organogenesis is dynamically regulated by constant signaling crosstalk between resident cells and their extracellular matrix (ECM) (Boudreau and Bissell, 1998) (Martins-Green and Bissell, 1995), (Silva et al., 2021), (Pally et al., 2022). Tissue specificity is fundamentally contingent upon extracellular cues that originate from, or are influenced by, the ECM (Xu et al., 2009). The core matrisome refers to the structural components of the ECM and consists of approximately 300 unique matrix macromolecules classified into collagens, proteoglycans (like heparan sulphate, hyaluronan), and glycoproteins (like laminins, elastin, fibronectin) (Naba et al., 2016). These ECM components are post-translationally modified by a range of secreted enzymes, like proteases, oxidases, and glycosidases. Remodeling of the ECM is a highly regulated physiological process occurring in development and in restoring tissue homeostasis during wound repair (Bonnans et al., 2014). It is not surprising that such processes are highly dysregulated in, and ECMs contribute etiologically to, pathologic conditions including cancer (Cox and Erler, 2011). In fact, the ECM may constitute as much as 60% of the tumor mass (Henke et al., 2020). Although cancer-associated fibroblasts (CAFs) are predominantly involved in secreting these ECM molecules, transformed tumor cells also contribute copiously to the construction of the tumor matrix microenvironment. Therefore, widespread genomic alterations in cancer cells may entail aberrations in ECM genes, that could in turn irreversibly alter their coded protein structure and function (Casey et al., 2009). Such alterations could adversely affect tissue architecture, disrupting homeostasis and spurring cancer progression.

A series of studies have investigated associations between ECM gene-level changes and tumorigenesis. For example, a frequent deletion of a region in chromosome 10 is associated with loss of heterozygosity, leading to aberrant expression of a tumor suppressor glycoprotein DMBT1 (mapped to that region) in lung cancer cells (Takeshita et al., 1999). Mutation of basement membrane protein FBN3 in small cell lung cancer cells is detected only in hilar lymph node metastasis but not in the primary tumor or other metastatic sites, suggesting that mutations may potentially steer progressions towards specific organotypic locales (Iwakawa et al., 2015). Similarly, genetic aberrations in laminin genes are frequently seen in gastric and colorectal cancers with high microsatellite instabilities (Choi et al., 2015). Although such studies underscore the fact that genetic instability associated with carcinogenesis could subsume ECM-encoding genes, systemic analyses on this topic have been missing. A notable exception is an elegant pan-cancer analysis that examined The Cancer Genome Atlas (TCGA) database to report that copy number alterations (CNAs) and mutations are more frequent in matrisomal genes than in the rest of the genome. These alterations were also predicted to significantly impact matrisomal gene expression and protein function. Moreover, a set of matrisomal genes were identified whose mutational burden could be used as an independent predictor of cancer survival (Izzi et al., 2020).

In this manuscript, we have focused on genomic alterations (mutations and copy number aberrations) in core matrisomal genes in nine cancers. We categorized the genes based on the pervasiveness of their mutations across cancers and their evolutionary age using phylostratigraphy. Biological interactions involving their encoded proteins, their interacting partners as well as their ontologically enriched functions were annotated. Our results reveal fascinating correlations between the pervasiveness of ECM glycoprotein mutation and their evolutionary age, which could explain the differential contributions of specific ECM glycoproteins to carcinogenesis.

## RESULTS

### Mutations are commonly seen in genes encoding the core matrisome across neoplasia

We sought to estimate the extent of aberrations specific to core matrisomal genes across a set of nine (breast, ovarian, prostate, colorectal, bladder, gastric, liver, lung, and skin) human cancers. These cancers represent transformations of epithelial cells but are diverse in their germ layer of origin: ectodermal (skin, breast), mesodermal (urothelium, ovarian, prostate), and endodermal (lung, liver, colorectal, and gastric); and their tissue histotype: skin (stratified squamous); breast, gastric, liver (cuboidal); urothelial (transitional); colorectal, prostate, lung (columnar); and ovarian (squamous-cuboidal or fallopian epithelia: cuboidal). Moreover, these cancers are among the leading causes of cancer-related deaths all over the world. We began by measuring the proportion of specific gene alterations (mutations, copy number amplifications, and deletions) across the nine cancers and further estimated the contribution of matrisome-encoding genes to each of these alterations. The proportion of genes undergoing copy number alterations was numerically higher than those showing mutations across all cancers (Supplementary File 1); this has also been observed for matrisome-encoding genes in general by Izzi and coworkers (Izzi et al., 2020). To our surprise, though, we observed that across all cancers, the proportion of core matrisome-encoding genes showing mutations (mean: 2.85, SD:1.28) was greater than those showing an amplification in their copy number (mean: 0.91, SD: 0.31) or deletion (mean: 0.61; SD: 0.26) (one way ANOVA, p < 0.001) (Figure 1A and B) (See also Supplementary File 1 and 2). Although cancers have been shown to belong to two distinct subsets, which show specific signatures of genomic aberrations due to mutations in one case and copy number variation in the other (Ciriello et al., 2013), our observations suggest specific number-limited gene sets may override such global trends and show similar signatures across the wide set of malignancies. In order to confirm whether such patterns of overrepresentation of mutation as a genomic alteration were seen in other related gene sets, we estimated the same in the list of matrisomal associated proteins (which include soluble protein factors, endogenous lectins and annexins, and proteases and their inhibitors). We observed no patterns: different sets of cancers showed a preponderance of mutational or copy number variational alterations (Supplementary Figure 1), consistent with the findings of Ciriello and coworkers (Ciriello et al., 2013).

**Figure 1.**
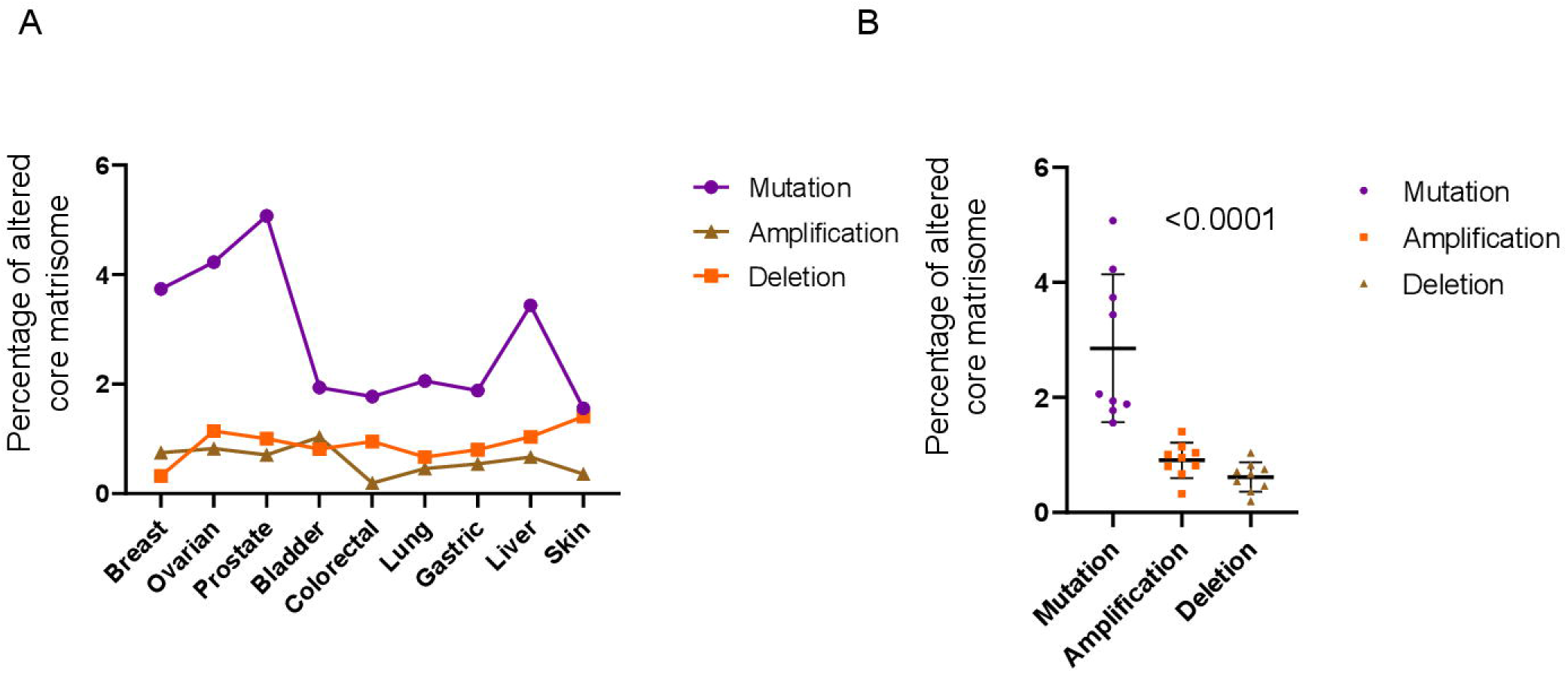
Frequency of mutations of the core matrisomal genes is dominant over amplification and deletion across nine human cancers. (A) The graph represents the core matrisomal genes as a fraction of the total number of human genes undergoing specific alterations across cancers. Purple points represent mutations, orange represents copy number amplifications, and brown represents ∞py number deletions. (B) Dot plot graph showing a comparison of the specific frequencies of 1A. Each dot corresponds to the frequency associated with a given cancer in our study. Statistical significance was computed using one way ANOVA with Tukey’s multiple ∞mparisons test for post hoc comparison. Bars represent mean+/- S.D. (See also Supplementary File 1 and 2).

The core matrisome has been classified biochemically into glycoproteins, proteoglycans, and collagens. We investigated the extent of alterations among the three groups of the core matrisomal genes. Of the mutated core matrisomal genes, glycoproteins constituted the predominant fraction (mean: 66% SD: 3.61) compared with proteoglycan- and collagen-encoding genes (mean: 9.5% SD: 2.40 and mean: 24.4% SD: 5.48, respectively) (Figure 2A, 2B and Supplementary File 1). Of the amplified core matrisomal genes, glycoproteins constituted the predominant fraction (mean 68.56 % SD: 4.04) compared with proteoglycan- and collagen-encoding genes (mean: 12.71 SD: 1.86 and mean: 18.71 SD: 3.86, respectively) (Figure 2C, 2D and Supplementary File 1). Of the deleted core matrisomal genes as well, glycoproteins constituted the predominant fraction (mean 76.20 % SD: 11.12) compared with proteoglycan- and collagen-encoding genes (mean: 9.72 SD: 9.72 and mean: 14.06 SD: 8.22, respectively) (Figure 2E, 2F and Supplementary File 1). While this was not surprising, given that the proportion of total genes encoding for glycoproteins exceeds the other two categories (and when normalized to the total number of genes within the subcategories, the collagen-encoding genes turned out to exhibit the biggest set with mutations), we nevertheless directed our attention for further studies to glycoproteins, given that this constituted the largest absolute set of mutations, whose effects on tissue architecture and carcinogenesis could be sensitively discerned (Figure 2A, C, and E).

**Figure 2.**
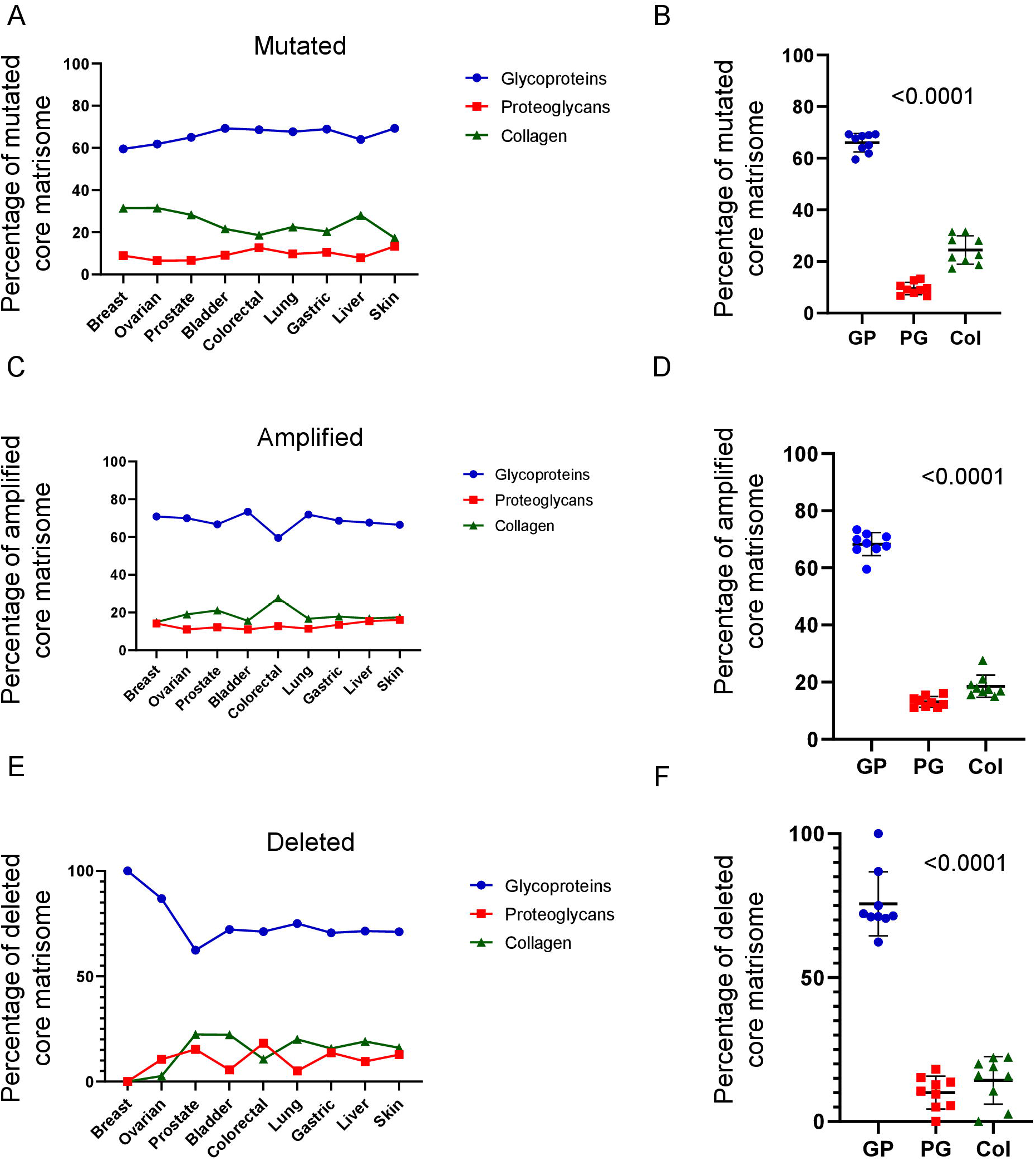
Matrisomal genes predominantly undergoing alterations in cancers are glycoproteins. (A) Graph representing the core matrisomal constituents: glycoproteins (GP, blue), proteoglycans (PG, red) and Collagens (Col, green) showing mutations as a percentage of mutated human genes across cancers.(B) Dot plot graph showing a comparison of the specific frequencies of 2A. (C) Graph representing the core matrisomal constituents: glycoproteins (GP, blue), proteoglycans (PG, red) and Collagens (Col, green) showing copy number amplifications as a percentage of amplified human genes across cancers. (D) Dot plot graph showing a comparison of the specific frequencies of 2C. (E) Graph representing the core matrisomal constituents: glycoproteins (GP, blue), proteoglycans (PG, red) and Collagens (Col, green) showing deletions as a percentage of deleted human genes across cancers. (F) Dot plot graph showing a comparison of the specific frequencies of 2E. Each dot corresponds to the frequency associated with a given cancer in our study. Statistical significance was computed using one way ANOVA with Tukey’s multiple comparisons test for post hoc comparison. Bars represent mean+/- SD. (See also Supplementary file 1 and 2).

We next investigated the ubiquity of mutations or copy number alterations for a set of glycoprotein genes across cancers. Based on the frequency of their alteration across tumors, we classified the glycoprotein genes into four subsets: 1. Universal (for which we found evidence of a specific alteration across all nine cancers); 2. Common (for which we found evidence of a specific alteration across seven to eight cancers); 3. Intermediate (for which we found evidence of a specific alteration across four to six cancers); and 4. Rare (for which we found evidence of a specific alteration across only one to three cancers) (see Table 1 for the mutated set of matrisomal glycoprotein genes, and Supplementary Tables 1 and 2 for the counterparts showing copy number amplification and deletion, respectively).

**Table 1:** Classification of genes encoding core matrisomal glycoproteins showing missense mutations across all nine cancers (denoted as Universal set), seven to eight of nine cancers (denoted as Common set), four to six of nine cancers (denoted as Intermediate set) and one to three of nine cancers (denoted as Rare set).

| Universal | Common |  | Intermediate |  |  | Rare |  |
| --- | --- | --- | --- | --- | --- | --- | --- |
| DMBT1 | ABI3BP | SLIT1 | AGRN | MEPE | VWA5A | BSPH1 | NTN4 |
| FBN1 | AEBP1 | SLIT2 | AMBN | MFAP1 | VWA5B1 | CTHRC1 | NTN5 |
| FBN2 | CILP | SLIT3 | BMPER | NDNF | VWA5B2 | EMID1 | PCOLCE |
| FBN3 | DSPP | TECTA | CILP2 | NTN1 | VWCE | IGFBP2 | RSPO1 |
| FN1 | EGFLAM | THBS1 | COCH | NTN3 | ZP1 | LRG1 | RSPO3 |
| FRAS1 | ELN | THBS2 | COMP | NTNG1 | ZP2 | MFAP3 | SPON2 |
| HMCN1 | EYS | TINAG | CRIM1 | OIT3 | NPNT | TINAGL1 | VTN |
| IGSF10 | FGA | TNC | CRISPLD1 | OTOL1 | DMP1 | RSPO4 | ZP3 |
| LAMA1 | FNDC1 | TNR | ECM1 | PAPLN | FGB | ADIPOQ | CRELD2 |
| LAMA2 | LAMA4 | VWA3B | ECM2 | PCOLCE2 | FGL2 | CDCP2 | IGFBP1 |
| LAMA3 | LAMA5 | VWA7 | EDIL3 | POSTN | IBSP | COLQ | MATN3 |
| LAMB1 | LAMB3 | VWDE | EFEMP1 | RSPO2 | LGI1 | CRELD1 | MFAP2 |
| LAMB2 | LAMC1 | ZP4 | EMILIN1 | SMOC2 | LGI2 | CRISPLD2 | POMZP3 |
| LAMB4 | LAMC2 |  | EMILIN2 | SNED1 | LGI3 | EFEMP2 | VWA1 |
| MXRA5 | LAMC3 |  | EMILIN3 | SPARCL1 | MATN1 | FBLN5 | IGFBP7 |
| PXDNL | LTBP1 |  | FBLN1 | SRPX2 | MMRN2 | FGL1 | AMELX |
| RELN | LTBP2 |  | FBLN2 | TECTB | NTNG2 | FNDC8 | ELSPBP1 |
| SVEP1 | LTBP4 |  | FBLN7 | THBS3 | SMOC1 | GLDN | KCP |
| TNN | MMRN1 |  | FGG | THBS4 | SPARC | IGFALS | DPT |
| TNXB | NELL1 |  | FNDC7 | THSD4 | SPP1 | IGFBP3 | IGFBP6 |
| VWF | NELL2 |  | GAS6 | TNFAIP6 | SRPX | IGFBP4 | IGFBPL1 |
|  | NID1 |  | HMCN2 | TSKU | TGFBI | IGFBP5 | MFAP4 |
|  | NID2 |  | LTBP3 | TSPEAR | VWA2 | LGI4 | MGP |
|  | OTOG |  | MATN2 | VIT | ZPLD1 | MFAP5 | SBSPON |
|  | PXDN |  | MATN4 | VWA3A |  | MFGE8 |  |

**Table 2:** Ontologically enriched (>1E-07) pathways associated with the universal and rare sets of mutated core matrisomal glycoproteins based on biological process

| Set | FDR adjusted p value | Functional Category |
| --- | --- | --- |
| Universal | 2.63E-09 | Cell adhesion |
|  | 2.63E-09 | Biological adhesion |
|  | 2.63E-09 | Extracellular matrix organization |
|  | 4.50E-09 | Extracellular structure organization |
|  | 2.07E-05 | Regulation of cell migration |
|  | 2.91E-05 | Regulation of cell motility |
|  | 4.49E-05 | Regulation of locomotion |
|  | 4.49E-05 | Regulation of cellular component movement |
| Rare | 3.45E-09 | Regulation of insulin-like growth factor receptor signaling pathway |
|  | 3.27E-08 | Insulin-like growth factor receptor signaling pathway |
|  | 1.25E-05 | Post-translational protein modification |
|  | 1.74E-05 | Regulation of canonical Wnt signaling pathway |

**Table 3:** Ontologically enriched (>1E-07) associated with the universal and rare sets of mutated core matrisomal glycoproteins based on cellular function

| Set | FDR adjusted p value | Functional Category |
| --- | --- | --- |
| Universal | 1.32E-23 | Extracellular matrix |
|  | 4.14E-19 | Collagen-containing extracellular matrix |
|  | 5.35E-16 | Basement membrane |
|  | 1.59E-13 | Extracellular region |
|  | 7.51E-12 | Extracellular region part |
|  | 2.33E-10 | Laminin complex |
|  | 5.10E-09 | Extracellular matrix component |
|  | 4.23E-06 | Laminin-3 complex |
| Rare | 6.92E-17 | Extracellular space |
|  | 4.99E-10 | Insulin-like growth factor binding protein complex |
|  | 4.99E-10 | Growth factor complex |
|  | 5.41E-09 | Endoplasmic reticulum lumen |
|  | 1.57E-07 | Insulin-like growth factor ternary complex |
|  | 2.94E-06 | Microfibril |

**Table 4:** Ontologically enriched (>1E-07) pathways associated with the universal and rare sets of mutated core matrisomal glycoproteins based on molecular function

| Set | FDR adjusted p value | Functional Category |
| --- | --- | --- |
| Universal | 5.30E-22 | Extracellular matrix structural constituent |
|  | 4.46E-13 | Structural molecule activity |
|  | 8.55E-07 | Integrin binding |
|  | 2.71E-05 | Cell adhesion molecule binding |
|  | 4.91E-05 | Signaling receptor binding |
| Rare | 1.83E-14 | Insulin-like growth factor II binding |
|  | 4.81E-13 | Insulin-like growth factor I binding |
|  | 1.71E-10 | Growth factor binding |
|  | 1.87E-08 | Heparin binding |
|  | 1.84E-07 | Glycosaminoglycan binding |
|  | 2.70E-07 | Sulfur compound binding |
|  | 8.33E-06 | Frizzled binding |
|  | 1.07E-05 | Signaling receptor binding |

### Universally mutated glycoprotein genes predominantly evolved from unicellular ancestors

We next sought to probe the reason behind such asymmetries in the ubiquity of glycoprotein genomic alterations. Trigos and colleagues have used phylostratigraphy to estimate the evolutionary age of genes based on the presence of gene orthologs within unicellular, early metazoan, and mammalian organisms’ genomes (Trigos et al., 2019). We employed a similar approach by segregating mutated, amplified, and deleted sets of glycoprotein genes into their respective phylostrata (Supplementary File 3).

To our surprise, a dominant fraction: 56.25% of the universally mutated glycoprotein gene set were predicted to be unicellular in origin, and 43.75% were predicted to have evolved in early metazoans (Figure 3A and Supplementary Files 2 & 3). In contrast, predominant fractions of common and intermediate mutated glycoprotein gene sets were early metazoan in their origin (64.86% and 82.86%, respectively), with relatively lesser fractions being specific to the unicellular-origin genes (35% and 13%, respectively), and even smaller fractions in the mammalian category (0% and 4.29%, respectively). The dominant fraction of the rare set of mutated glycoprotein genes were predicted to have evolved with metazoans (84%), 14% evolved in mammals, and 2.3% evolved in early unicellular organisms. Within the amplified glycoprotein genes, the dominant fraction of universal, common, intermediate, and rare sets evolved in early metazoans (57%, 67%, 79%, and 77%, respectively), with a smaller proportion being unicellular in origin (42%, 32%, 12%, and 14%, respectively), and with a minor representative set of genes only in the intermediate and rare set (8% and 7%) having evolved in mammals (Figure 3B and Supplementary Files 2 & 3). The universal and common set of ECM glycoprotein genes that showed deletion were entirely early metazoan in origin (which can be explained by the fact that this subset consisted of only 2 genes). The intermediate and rare sets were predominantly unicellular in origin (53%, 15%, respectively), and early metazoan in origin (40%, 79%, respectively), with small fractions being mammalian origin (7%, 6%, respectively) (Figure 3C and Supplementary Files 2 & 3).

**Figure 3.**
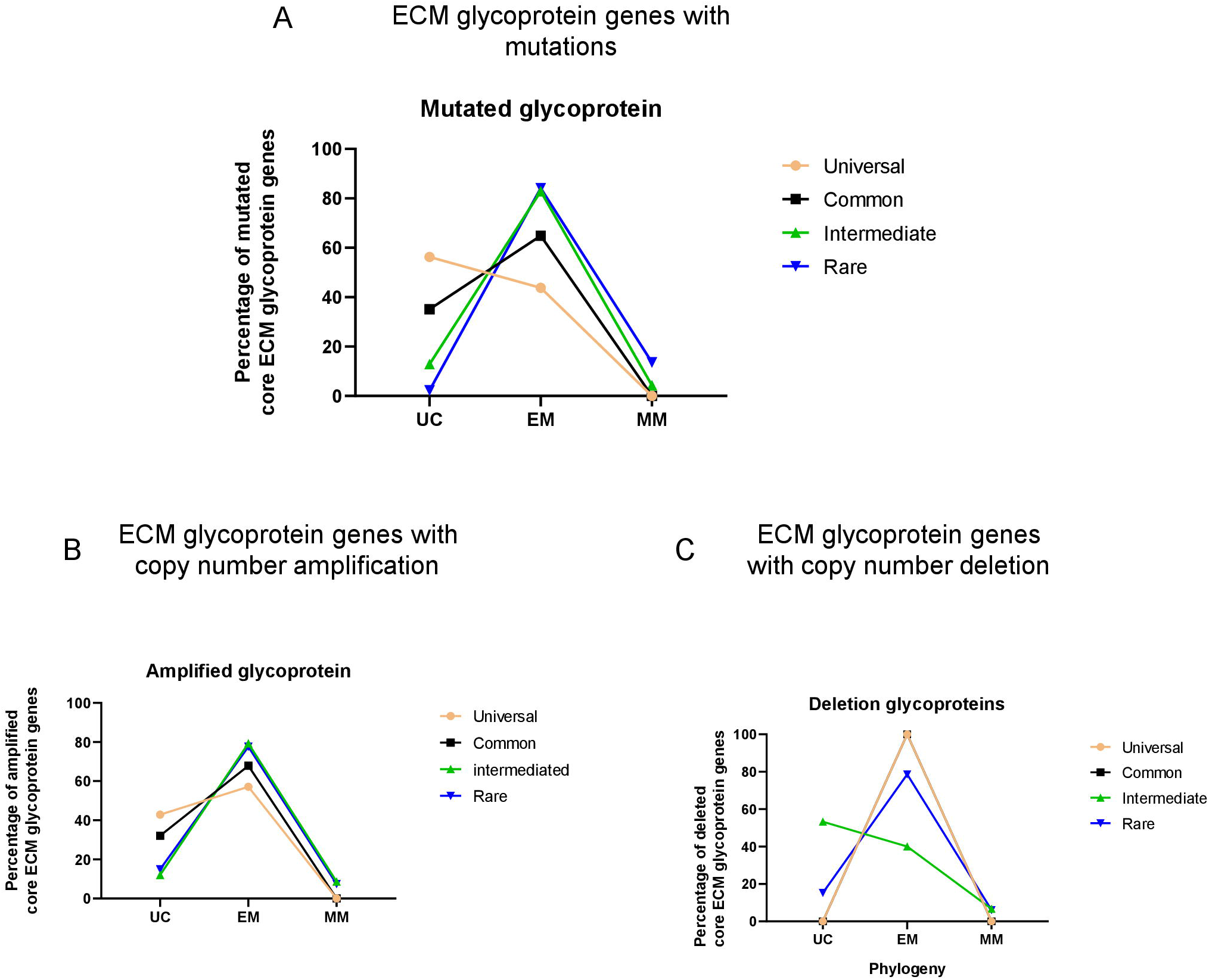
Association between gene age and frequency of genomic aberration across cancers for core matrisomal glycoproteins. Graphs represent the percentage of core matrisomal glycoprotein genes showing (A) mutations, (B) copy number amplification and (C) deletions as a function of their gene age measured through phylostratigraphy (phylostrata 1 to 3 represent unicellular origin, phylostrata 4 to 9 representing early metazoan origin while phylostrata 10 to 16 represent mammalian-specific origin of genes) Genes are also arranged based on the extent of their aberrations across nine cancers (peach: universal set; aberrations across all nine examined cancers, black: common set; aberrations in 7-8 of nine cancers, green: intermediate set; aberrations in 4-6 of nine cancers, blue: rare set; aberrations in 1-3 of nine cancers.) (See also Supplementary File 3).

The predominant type of genetic alteration was probed for in the two extreme subsets: universal and rare. In both cases, missense mutations showed the highest frequency with other alterations such as nonsense and frameshift mutations, splicing, and fusion bearing minor proportions (Supplementary Figure 2). In addition, the missense mutations were further analysed for the preponderance to deleteriously affect protein function using two independent algorithms: SIFT (Sorting Intolerant from Tolerant, analyzing the consequences of amino acid replacement on protein function based on primary structure and biophysical characteristics of residues) (Ng and Henikoff, 2001), and the PolyPhen-2 algorithm (which examines the consequences of amino acid replacement by looking at changes at the level of both sequence and predicted structure and uses a Naïve Bayes classifier to arrive at an interpretation of the effect on function (Adzhubei et al., 2010). We discovered that in the universal gene data set, the proportion of “SIFT-deleterious” and “SIFT-tolerated” mutations are higher when compared with its rare counterpart. The former is potentially associated with predicted loss-of-function mutations, whereas the latter may bear an overlap with mutations resulting in gain-of-function. Similarly, the PolyPhen-2 algorithm showed a greater propensity for both benign effects (overlapping with gain-of-function) as well as damaging effects (loss of function) in the universal set compared with the rare set, suggesting a greater tendency for cancer-pervasive mutations to perturb functions of ECM glycoprotein.

### Universally mutated ancient glycoproteins regulate a greater degree of molecular reactions

We next sought to probe whether variations in the ubiquity of the universal and rare set of mutated glycoprotein genes could be explained by differences in the ability of their coded proteins to regulate or be regulated by other proteins. To do so, we annotated regulatory interactions from PathwayCommons.org for every gene from the universal and rare sets and performed network analysis to identify upstream genes that control the gene of interest and downstream genes that are controlled by the gene of interest. Although there was no significant difference in the extent to which the universal and rare mutated ECM glycoprotein sets were regulated (Figure 4B and Supplementary Files 2 & 4), the mean proportion of genes, whose expression was regulated in turn by the universal set of ECM glycoproteins were found to be significantly higher than that of the rare set (Figure 4C and Supplementary Files 2 & 4). This implies that a disruption in activity due to mutations of the universal set of glycoproteins could potentially disrupt stable states within larger networks of proteins and the cellular reactions they are involved in.

**Figure 4:**
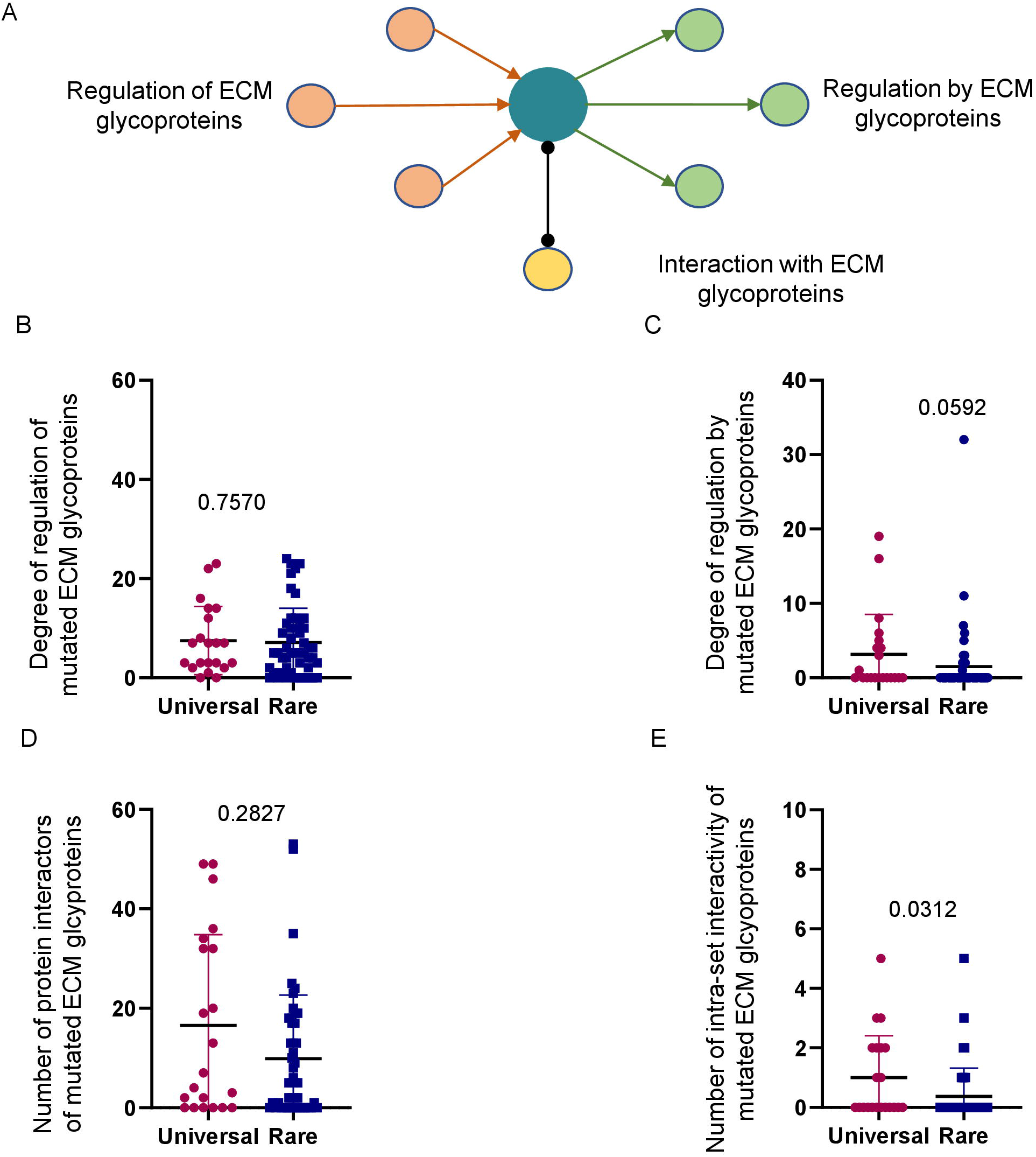
ECM glycoproteins mutated across most cancers tend to regulate and interact within the set with a wider set of proteins than those mutated across fewer and specific cancers. (A) Schematic depicting estimations of reactions that regulated, or in turn were regulated by, a given ECM glycoprotein as well as the number of interactions with other proteins. (B) Graph representing the percentage of the upstream genes that regulate the universal and rare set of core matrisomal glycoproteins. (C) Graph representing the percentage of the downstream genes that are regulated by the universal and rare set of core matrisomal glycoproteins. Each dot represents the number of genes upstream or downstream of a particular member in the universal (magenta)/rare (blue) gene set. (D) Graph representing the number of protein interactors for each gene present in universal or rare set of core matrisomal glycoproteins. Each dot represents the number of interactors for a given gene. (E) Graph representing the number of intra-set interactors within the universal or rare set of core matrisomal glycoproteins. Each dot represents percentage of interactors for a given gene that are members of the set from which gene is taken. Statistical analysis was performed using unpaired, non-parametric test. Bars represent mean +/- SD (See also Supplementary File 4 and 5)

Furthermore, we compared the number of elucidated interacting proteins for each of the mutated ECM glycoproteins in the universal and rare sets. Interestingly, the average number of interactors for ECM glycoproteins belonging to the universal and rare sets showed no difference. (Figure 4D and Supplementary Files 2 & 5). However, the proteins encoded by the universal set showed a greater tendency to interact with each other than was observed between their rare set counterparts (Figure 4E and Supplementary Files 2 & 5). These observations suggest that mutations in the universal ECM glycoprotein set may have more detrimental consequences regarding disruptions of protein-protein interactions and dysregulation of downstream genes.

### The ECM glycoproteins of the universal set regulate fundamental organogenetic functions

We next investigated the ontological associations of the universal and rare set of ECM glycoprotein genes showing mutations. Comparing the top ontological categories revealed specific enrichments for the two sets. The distinct biological processes for the mutated universal glycoprotein gene set were cell adhesion, regulation of basement membrane organization, and cell-substrate adhesion, whereas those for the rare set were regulation of insulin-like growth factor receptor signaling pathway, extracellular structure organization, external encapsulating structure organization, and developmental process (Supplementary Files 6 and 9). Similar results were seen in the case of enrichment using molecular functions (Supplementary files 8 and 11) and cellular processes (Supplementary files 7 & 10): the rare set was more associated with extracellular signaling, whereas the universal set correlated with morphogenesis and associated phenomena such as adhesion and integrin-based signaling, reinforcing the notion that their disruption through mutations is likely to disrupt conserved tissue homeostatic programs.

## Discussion

The sheer number of genetic aberrations seen within and across transformed epithelia in a tumor renders it difficult to ascertain the change in the activity of which gene product(s) is causative to malignant dedifferentiation and dysregulation of phenotypic behaviors associated with cancer (Martincorena et al., 2017), (Stratton et al., 2009). On the other hand, while most of such aberrations may only be consequential to the genomic instability brought about by a few “driver” mutations (although Dressler and colleagues’ (Dressler et al., 2022) compendial approach suggests a large number of canonical and candidate drivers of cancer), the former may nevertheless influence the tropism of the tumor. Whichever the case may be, one approach to investigating the mechanisms of loss of tissue architecture, a hallmark of all cancers (Bhat and Bissell, 2014), could be to focus on gene products responsible for establishing the architecture in the first place. Therefore, it is unsurprising that the disruption in functions of the extracellular matrix proteins within oncological microenvironments has received so much attention in the last few decades (Pickup et al., 2014), (Lu et al., 2012).

While the regulation of ECM organization through proteolysis, aberrant synthesis, and reorganization have provided exciting insights into how cancers grow, persist, and debilitate (Winkler et al., 2020), whether there are specific patterns in aberrations of ECM genes associated with cancer, and if so, to what extent, has received relatively scant attention. In this manuscript, we have taken on these questions to identify a set of ECM glycoproteins, which commonly show missense mutations across a set of nine cancers. The diversity in the embryological origin and histology of these cancers suggests that the disruption in functions of these genes may affect morphogenetic mechanisms central to distinct organogenetic programs.

When the ECM glycoprotein set was ontologically assessed, it was unsurprising that the molecular functions that were statistically enriched were associated with the formation of the basement membrane, one of the canonical mediators of glandular morphogenesis and homeostasis (Pozzi et al., 2017). Consistent with ontological predictions, the set consists of various members of laminin- and tenascin family of genes, which code for principal basement membrane proteins (Pozzi et al., 2017) (Akimoto et al., 1992). Another enriched family is that of fibrillins, ECM proteins that constitute stromal microfibrils closely interacting with BM structures (Tiedemann et al., 2005) (Eckersley et al., 2018). Other genes encode proteins associated with cell migration (RELN, FN1, FNDC1) (Yuan et al., 2012) (Hsiao et al., 2017) (Liu et al., 2019), and TGF-β activity (LTBP1, MXRA5) (Tritschler et al., 2009) (Poveda et al., 2017), whose function in morphogenesis is less well understood. Given our results, it is unsurprising that as much as 57% of ECM glycoproteins that are mutated across cancers are candidate drivers of carcinogenesis compared with 6% of glycoproteins that are mutated specifically across one or two cancers.

A predominant proportion of these genes have orthologs with unicellular organisms. A more global analysis of genomic aberrations associated with human cancers led investigators to suggest that the genes which showed the highest frequencies of point mutations and copy number amplifications, originated in early metazoan genomes (Trigos et al., 2019). Another study predicts the emergence of gatekeeper genes (that are involved with tissue homeostasis) during early metazoan evolution, whereas caretaker genes (involved in genomic stability) emerged within unicellular organisms (Domazet-Loso and Tautz, 2010). Cell biological investigations, however, show that the signaling cues driven through homeostatic ECM scaffolds regulate genomic stability within tissues suggesting such deep unicellular links between phenotype (ECM-cell architecture) and genotype (in terms of genomic fidelity) (Radisky and Bissell, 2006) (Sonugür and Akbulut, 2019). In contrast, ECM glycoproteins that were mutated across fewer cancers showed a greater predisposition towards evolutionary innovation in metazoans and even mammals. Our network analysis implies that a longer evolutionary time may have provided greater opportunities for ancient ECM glycoproteins to form larger networks wherein they integrated better with each other to regulate a larger set of transductive reactions within cells. This allowed them to influence a larger set of morphogenetic programs cutting across histological constraints. Our work is also consistent with theoretical explorations of cancer as an atavistic reversion to unicellular modes of phenotypic behavior, as has been formalized by Davies and colleagues (Lineweaver et al., 2021), (Bussey and Davies, 2021). Their suggestion for the use of phylostratigraphy bears out in our effort that uses similar approaches to showcase the correlation of age of innovation of ECM proteins with the cancer-pervasive effects of their mutations.

Our analysis is not without its limitations. We chose a relatively higher cut-off frequency (greater and equal to 1.0%) for point mutations that allowed us to enrich an optimal set of glycoproteins with specific ontological and interactive patterns. However, the same threshold may have resulted in fewer putative targets that showed copy number variation. Secondly, despite the prolific use of phylostratigraphy as a means of ascertaining the evolutionary origin of genes (Holland, 2015), (Domazet-Lošo et al., 2017), (Zhang et al., 2019), (Domazet-Lošo et al., 2007), the method has received criticism for not considering heterogeneities in rates of gene evolution across phylogenetic lineages (Moyers and Zhang, 2017). Notwithstanding such drawbacks, our observations provide a quantitative understanding of the extent to which the dysregulation of the core matrisomal genome associates across the oncological paradigm. In the future, we would seek to search deeper within the core matrisomal sets to identify patterns within copy number variations as well as extend our analyses to genes encoding proteins that regulate the remodeling of the extracellular matrix.

## Materials & Methods

### Genomic Alteration Data

The lists of mutated and copy number altered (amplified and deep deleted) genes for nine cancer types (Breast, Ovarian, Prostate, Bladder, Colorectal, Gastric, Lung, Liver, Skin) were obtained from the cBioPortal web server (https://www.cbioportal.org/) (Cerami et al., 2012) (Gao et al., 2013). Under the ‘Query’ tab on the cBioPortal homepage, each cancer type was looked up either using the left-hand panel or the top-right search box (e.g., Breast, Ovary/Fallopian Tube, etc.). After checking the box ‘Select all listed studies matching filter’, any studies involving metastatic tumours, xenografts, etc., were deselected. With the remaining studies checked, ‘Explore Selected Studies’ was clicked. On the next page, under the ‘Summary’ tab, the data of ‘Mutated Genes’ and ‘CNA Genes’ was downloaded from the respective information boxes. (For list of studies chosen for this investigation, see also Supplementary Table 9)

### Core Matrisome Gene List

The gene list of core matrisome genes, i.e., glycoproteins (GP), proteoglycans (PG), and collagens (Col) in humans, was obtained from the Matrisome Project web server (http://matrisomeproject.mit.edu/) (Naba et al., 2016). From the homepage, after proceeding with ‘Human Matrisome’ under the ‘*In silico* Matrisome’ tab, the Excel files (.xls) were separately downloaded for ECM glycoproteins, collagens, and proteoglycans under the ‘Core Matrisome’ heading.

### Data Parsing for Downstream Analysis

A Python script written in-house was employed to isolate the list of core matrisomal genes that are altered in nine different types of cancers. The set of altered (mutated and copy number aberrated) matrisomal genes that had a frequency of alteration less than 1% (i.e., percentage of samples with one or more alterations in a particular cancer) was eliminated and proceeded with the set of altered genes having a frequency of alteration greater than equal to 1% for each of the nine cancers.

### Categorization according to Pervasiveness of Alteration

The copy number aberrated core matrisome genes were divided into amplified and deep deleted lists. The set of altered genes was compared across the nine cancers, and the genes were categorized according to their pervasiveness of alteration, viz. universal (altered across all nine cancer types), common (altered in seven to eight cancers), intermediate (altered in four to six cancers) and rare (altered in one to three cancers). The fraction of altered matrisomal genes across the different phylostrata in each of the 4 categories, i.e., universal, common, intermediate, and rare, was plotted.

### Phylostratigraphy of Core Matrisome Genes

The evolutionary ages of genes classified (using phylostratigraphy) into a range from 1 to 16 were acquired from previously published literature (Trigos et al., 2017) (Domazet-Lošo et al., 2007), where phylostrata 1 to 3 represent unicellular ancestors (UC), phylostrata 4 to 9 represent early metazoans (EM) while phylostrata 10 to 16 represent mammalian-specific genes (MM). Using another in-house Python script, the list of altered (mutated and copy number aberrated) matrisomal genes was matched against their respective phylostrata.

### Access and Categorization of Mutational Information

To investigate the types of mutations and their distribution across different cancers, the studies pertaining to a particular cancer are selected (as per the method described above), only ‘Mutations’ is checked next to the ‘Select Molecular Profiles’ section, the list of Universal or Rare altered genes was copy-pasted in ‘Enter Genes’ box under the ‘User-defined list’ drop-down, and ‘Submit Query’ was clicked on. On the next page, under the ‘Mutations’ tab, each gene from the list was individually selected, and information on mutations was downloaded by clicking on the ‘Download TSV’ button. Finally, the different types of mutations (missense, nonsense, fusion, etc.) were analysed for mutated glycoproteins across nine different cancers.

### Gene Regulatory Network

A human gene regulatory network (GRN) from the Pathway Commons database (https://www.pathwaycommons.org/) (Cerami et al., 2011) (Rodchenkov et al., 2020) was obtained for every altered gene in the universal and rare list of altered glycoproteins by obtaining the set of interactions connected by edges of the type ‘control-expression-of’ and ‘control-of-state-change’. While the number of outgoing edges of a gene represents the extent of its downstream regulatory network, the number of incoming edges of a gene is proportional to the degree of being regulated. The degree of regulation by and of a particular altered ECM gene was calculated for every gene in the universal and rare sets of altered glycoproteins.

### Protein-Protein Interaction

The set of protein-protein interactions (PPI) was obtained for every altered glycoprotein in the universal and rare sets from the inBio Discover server (https://inbio-discover.com/) (Li et al., 2017). For every gene in the universal set, the number of interactors was calculated, and the distribution was compared against that of the interactors for the rare set. Finally, the proportion of interactors from a particular set to the members of that same set (e.g., the fraction of interactors for a universally altered gene that are part of the universal set) was determined.

### Gene Ontology analysis

Gene ontology analysis for universal and rare genes was performed using ShinyGO v0.61 graphical tool for gene enrichment analysis using the default setting for p-value cut off by selecting species human.

## Acknowledgements

This work was supported by the Wellcome Trust/DBT India Alliance Fellowship grant (IA/I/17/2/503312) awarded to R Bhat. R Bhat would additionally acknowledge support from the Department of Biotechnology, India (DBT) (BT/909 PR26526/GET/119/92/2017), and the Institute of Eminence grant (IE/CARE-19-0319). J Langthasa is supported by Senior Research Fellowship (SRF) from the Ministry of Education, India. S Mishra acknowledges the Prime Minister Research Fellowship support for his graduate research. Monica U acknowledges the KVPY program for scholarship support. R Bhat also benefited from discussions with members of the ‘Cellular agency in multicellular development and cancer’ cluster that is part of the Agency, Directionality and Function cohort program, funded by the John Templeton Foundation.

